# Morpho-molecular mechanisms study of photodynamic therapy and curcumin larvicide action on wild mosquitoes larvae of genus *Aedes*

**DOI:** 10.1101/2020.07.13.200295

**Authors:** BP Araújo, EA Silva, LP Rosa, NM Inada, I Iermak, RA Romano, NF Mezzacappo, FF Melo, FC Silva, MP Rocha, RAA Silva, MPL Galantini, EA Silveira, M Garbuio

## Abstract

**Introduction:** Until the first two weeks of October 2019, 1.489,457 probable dengue cases have been reported in Brazil, with an incidence rate of 708.8 cases per 100 thousand inhabitants. Still in 2019, in the same period, 123.407 probable cases of chikungunya were reported, with 15 deaths confirmed by clinical and epidemiological criteria. Regarding Zika, in that period, 10.441 probable cases were recorded, been the northeast region with the highest number of notifications, followed by the midwest one. It is well known that current policies to control the vectors of those arboviruses are not effective. Studies for use of light-activated photosensitizers as an alternative to conventional insecticides for sustainable control of mosquitoes vector such as *Aedes* (dengue, yellow fever, chikungunya, zika), *Anopheles* (malaria), *Culex* (yellow fever) can already be found showing advantages over conventional insecticides (efficacy, safety, non-mutagenicity and fast degradation).

**Objective:** To evaluate the effectiveness of Photodynamic Therapy (PDT), mediated by curcumin and blue LED (460nm) in mortality of wild mosquitoes larvae of genus *Aedes* and also to verify, through confocal microscopy, how the photosensitizer internalizes in larvae organism. In addition to evaluating the action of PDT on the larvae with Raman spectroscopy and histological technique.

**Materials and methods:** Ovitraps were placed in the city of Vitória da Conquista, Bahia, and the larvae collected in stages L2 and L3 were fed for 24 hours with curcumin in concentrations of 10, 20 and 50% mixed with fish feed and then subjected to irradiation with blue LED for 2h (22mW / cm^2^ and 158.4 J / cm^2^). The larvae were placed in a container with dechlorinated water and mortality was monitored for 24 and 48 hours. Control groups in which only the larvae were exposed to blue LED for 2 hours and in which the larvae were only fed with fish feed mixed with curcumin (10, 20 and 50%) were included in the study, in addition to the group without treatment. All experiments were repeated after a 2-month interval to confirm the results, totaling 240 tests (tests 1 and 2, n = 15) between groups PDT 10%, PDT 20%, PDT 50%, curcumin 10%, curcumin 20%, curcumin 50%, blue LED and untreated group. The larvae belonging to the PDT 20% group, 20% curcumin, blue LED and control were submitted to histological slides, confocal microscopy and Raman spectroscopy. Larvae mortality rates were compared between groups using univariate descriptive analysis.

**Results:** All PDT groups showed larvicidal activity, with the PDT group 20% showing the highest larval mortality in the shortest time. The images from confocal microscopy by laser scanning showed that curcumin was distributed throughout the digestive system of larvae and the analysis by Raman microspectroscopy have shown patterns of alteration and cell death, corroborated by histological sections.

**Conclusion:** It was concluded that PDT in all concentrations was effective in larval mortality, with PDT 20% having the best activity with mortality of 100% in 24 hours.

## INTRODUCTION

In recent years, the world has been challenged by the growing of diseases known as arboviruses. About 4 billion people live in areas at risk for transmission of dengue virus [1]. Dengue is the main arbovirus transmitted by *Aedes* mosquitoes, the main ones being *A. aegypti* and *A. albopictus*. It is estimated that 390 million new cases emerge every year, of which 500 thousand evolve to more severe cases that require hospitalization with more than 20 thousand deaths, the majority in tropical countries [2].

Recently, the growth of diseases such as chikungunya and zika has attracted the interest of the international scientific community to tackle arboviruses, due to the increasing incidence, expansion of geographic reach, possible effects caused by co-circulation of viruses and high costs for health system [3]. In Brazil, dengue virus reemerged in the 1980s and, since then, has been responsible for successive epidemics. Currently, about 90% of Brazilian municipalities have been infested by *A. aegypti*, also favoring the growth of cases of chikungunya and zika [4].

Zika virus was detected in Brazil in 2015 and it has the ability to spread through non-vector methods, such as vertical transmission, sexual contact and blood transfusion. In addition, consequences such as Guilláin-Barré Syndrome and microcephaly in babies whose mothers were affected by the virus during pregnancy, increase the severity of the problem [3]. Chikungunya fever, a disease that appeared in national territory in 2014, is associated with the possibility of persistent clinical manifestations in the chronic phase, which negatively interferes with patients’ quality of life [4].

Prevention and control activities for arboviruses in the country have been based on the work of agents to combat endemics and Community health agents. Those professionals work to raise the population’s awareness of the importance of controlling mosquito breeding sites and eliminating those vectors [5]. However, insecticides play a very important role in control actions in the sense of being widely used, both from the point of view of public managers and population in general [6].

New tools have been developed to act to combat the insect vector, in view of the growing resistance to insecticides by insects, in addition to the possible environmental and health impacts of people. The technique of sterile insects and use of bacterial and fungal agents stand out [1]. The use of photosensitisers as an alternative to conventional insecticides has also been studied. Such compounds are highly effective and safe, paving the way for photodynamic therapy (PDT) to become a possibility as a vector control technique [7; 9; 14].

PDT consists of combining a light source with a specific wavelength or within a certain range with a chemical compound known as photosensitiser, which in presence of molecular oxygen, promote the destruction of a biological target [8]. Studies with *Curcuma longa* have demonstrated its great potential as a photosensitiser in PDT, as well as its essential oils and natural derivatives, for the control of *Aedes* mosquito larvae [9].

Therefore, the objective of this study was to evaluate the effectiveness of PDT mediated by curcumin at concentrations of 10%, 20% and 50% associated with blue LED (460nm) on mortality of *Aedes* mosquitoes wild larvae. We also aimed to verify how the photosensitizer internalizes in the larvae organism through laser scanning confocal microscopy technique, in addition to evaluating the action of PDT on the larvae with Raman microspectroscopy and histopathological analyzes.

## MATERIALS AND METHODS

### Collection of material

This study was carried out in the municipality of Vitória da Conquista, Bahia, Brazil. Ovitraps have been strategically distributed throughout the municipality according to the areas of activity of community health agents. Eucatex palettes (6×15cm) were placed weekly in those traps, and after 72 hours they were removed, wrapped in newsprint and transported to Medical Entomology Laboratory at UFBA / CEMAE, located at Family Health Training School, in that municipality.

### Obtaining the larvae

In laboratory, palettes containing mosquito eggs were placed in a number of 3 or 4 in 5.5L plastic trays containing 500mL of dechlorinated water, in order to promote hatching eggs.

The trays were inspected daily for a period of up to 7 days to check for the presence of larvae. The collected larvae were then transferred to another tray with 500mL of dechlorinated water and fed with Goldfish Color Alcon® fish food (Camboriú, SC, Brazil) until they reached maturation stages 2 or 3.

### Preparation of diets containing curcumin

The amount of 100g of Goldfish Color Alcon® fish feed (Camboriú, SC, Brazil) was ground to obtain a fine and homogeneous powder and from this powder curcumin photosensitizer (Sigma-Aldrich, ref. C1386) was incorporated, obtaining mixtures of feed with 10%, 20% and 50% curcumin. Those mixtures were stored in dark flasks, wrapped in aluminum foil, and kept refrigerated.

### Distribution of larvae among experimental groups

240 tests were carried out whose larvae were distributed among 8 experimental groups. Control group have not undergone any treatment (P-L-); blue LED group was the one in which the larvae were only submitted to blue LED (P-L +); 10% curcumin group (P + L- / 10%) was the one in which the larvae were only fed with a ration containing 10% photosensitizer; curcumin group 20% (P + L- / 20%) was the one in which the larvae were only fed with a ration containing 20% photosensitizer; curcumin group 50% (P + L- / 50%) was the one in which the larvae were only fed with ration containing 50% of photosensitizer; PDT 10% group (P + L + / 10%) was the one in which the larvae were fed with a ration containing 10% photosensitizer and submitted to blue LED; PDT 20% group (P + L + / 20%) was the one in which the larvae were fed with ration containing 20% of photosensitizer and submitted to blue LED and PDT 50% group (P + L + / 50%) was the one in which the larvae were fed with ration containing 50% of photosensitizer and submitted to blue LED. The experimental procedures were performed in duplicate (test 1 and test 2) with an interval of 3 months between them (June and September), as shown in table 1.

**Table 1:** Distribution of larvae among experimental groups

| **GROUP** | **TEST 1**<br>**n** | **TEST 2**<br>**n** | **TOTAL** |
| --- | --- | --- | --- |
| **P-L-** | 15 | 15 | 30 |
| **P-L+** | 15 | 15 | 30 |
| **P+L-/10%** | 15 | 15 | 30 |
| **P+L-/20%** | 15 | 15 | 30 |
| **P+L-/50%** | 15 | 15 | 30 |
| **P+L+/10%** | 15 | 15 | 30 |
| **P+L+/20%** | 15 | 15 | 30 |
| **P+L+/50%** | 15 | 15 | 30 |
| **TOTAL** | 120 | 120 | 240 |

P= Photosensitizer/L= Led Light

### Experimental Procedures

To carry out the study, the larvae that reached maturation stages 2 or 3 were transferred to new plastic tray containing 500mL of dechlorinated water, where they were fed a total of 200mg of ration (without photosensitizer or with photosensitizer in concentrations of 10%, 20% and 50% depending on the experimental group) for a period of 24 hours. The trays were covered with perforated fabric and left in a dark environment to reduce the incidence of light.

In test groups to observe the effectiveness of photosensitizer curcumin 10%, 20% and 50% (P + L- / 10%, P + L- / 20%, P + L- / 50%), those larvae were monitored by 24h and 48h period where mortality was observed.

To carry out the tests of PDT groups (P + L + / 10%, P + L + / 20%, P + L + / 50%), after the 24h feeding period with the mixture of ration and photosensitizer, the larvae were transferred for 24 well cell culture plate wells containing 1.5mL of sterile dechlorinated water in each well. The plates were adapted to Biotable® RGB equipment (MMOptics, São Carlos, SP, Brazil) and were subjected to the maximum intensity irradiation cycle with blue LED for 2 hours (22mW / cm^2^, 158.4J / cm^2^). In experimental groups in which the effectiveness of LED light used individually was observed (P-L +), the larvae were not previously fed with mixtures of ration and photosensitizer, having been directly transferred to be subjected to LED irradiation phase in the equipment Biotable® RGB. The larvae have been then monitored for 24h to 48h to check for mortality.

### Histopathological Analysis

For preparation of histological slides, larvae which were fed with diet containing curcumin in the proportion of 20% and irradiated with blue LED under the same experimental conditions as the mortality tests (P + L + / 20%) were used. Immediately after irradiation, larvae still in a lethargic state were taken to fix in 1mL of 10% formaldehyde inside plastic ependorfs. Larvae that have not undergone any treatment (P-L-), which have only been subjected to blue LED (P-L +) and that have only previously been fed with a 20% ration and photosensitizer (P + L- / 20%) were also taken for fixation.

Then, larvae were submitted to a dehydration process, being placed one by one in plastic cassettes and dipped in alcohol in increasing concentrations (70%, 80%, 90% and 100%) for a period of 30 minutes in each one. Subsequently, they were dipped in xylol for a period of 30 minutes, for the alcohol removal process, allowing inclusion with paraffin.

The larvae were then dipped in paraffin for 1 hour to start the inclusion process, having been dipped again in another beaker also containing paraffin for another 1 hour. After inclusion, they were then inserted into previously heated metal molds containing paraffin for packaging. After cooling, paraffin blocks were removed from molds and taken to microtome MRP 2016® (Lupetec, São Carlos, Brazil) to perform cuts at 4 µm. The cuts obtained were then placed on slides that dried in an oven at 60°C to remove the paraffin and later stained with hematoxylin-eosin for visualization.

### Confocal Laser Scanning Microscopy

In analysis of confocal microscopy, the larvae of group P + L- / 20% and control (P-L-) were sent to sector of confocal microscopy, properly covered with aluminum foil to avoid previous exposure to light, and analyzed with excitation of 458nm. The larvae of control group (P-L-) were also analyzed at 488 nm, to demonstrate in this wavelength they have a little autofluorescence. The fluorescence of larvae associated with curcumin were sufficient to generate images that would make it possible to visualize the distribution of photosensitizer by the larval digestive system. The images were obtained using a Zeiss LSM780 inverted confocal fluorescence microscope (Carl Zeiss, Jena, Germany), using 10X objective lenses and using the Zen 2010 software (Carl Zeiss, Jena, Germany).

### Raman microspectroscopy

Larvae intestines were placed on glass slide of microscope covered with aluminum to remove fluorescence from the glass. Raman spectra were taken from the larvae intestinal tissues after each treatment. Larvae that did not undergo any treatment (P-L-) were analyzed, that only fed with the mixture of ration and photosensitizer at 20% (P + L- / 20%) and that were submitted to PDT after feeding with the mixture with 20% photosensitizer (P + L + / 20%).

For Raman microspectroscopy analyses, measurements were performed using a WITec Alpha 300 RAS microscope (WITec, Ulm, Germany). Excitation wavelength was 785nm and detection range was 100-3200 cm^-1^. Spectra were collected with a 20x magnification objective (Zeiss, Jena, Germany) and recorded with an integration time of 120s and 5 accumulations per spectrum. The obtained spectra were processed using WITec ProjectFOUR software and Origin 2016 software.

### Statistical analysis

To evaluate the effectiveness of different protocols tested for larval mortality, GraphPad Prisma 5 program (GraphPad Software, La Jolla, CA, USA) was used, and data were submitted to a univariable descriptive analysis to compare the rates of mortality among the groups tested at proposed times.

## RESULTS

### Larval mortality after procedures

PDT proved to be effective in eliminating the larvae in all tested parameters, been, PDT 20% group, able to eliminate all larvae after 24 hours of the procedure.

Larval mortality data in different groups were compared after 24h and 48h. After 24 hours, in both test 1 and test 2, it was possible to observe that in control groups (P-L-), the number of larvae remained the same as the initial one. In LED (P-L +) and 10% curcumin (P + L- / 10%) groups, there were a mortality rate of around 2 larvae, while in the other curcumin (P+ L- / 20% and 50%) groups, mortality were 2 to 6 larvae in both tests. In PDT groups (P + L + / 10%, 20% and 50%) there were deaths of more than 12 larvae from the 15 tested, and in that 24h time, PDT group 20% (P + L+ / 20%) reached the total elimination of all 15 tested, both in test 1 and 2. At 48h, in control groups (P-L-), the number of larvae also remained the same as in the initial group and in LED groups (P-L +) and in curcumin groups (P + L- / 10%, 20% and 50 %) more than 2 to 3 larvae died in relation to the number observed within 24 hours. In PDT groups (P + L + / 10%, 20% and 50%), only in test 1 PDT 10% group (P+ L + / 10%) has eliminated all larvae, while the other groups eliminated an additional 2 to 3 larvae in relation to the number observed at 24h, as shown in figures 1 and 2.

**Fig. 1.**
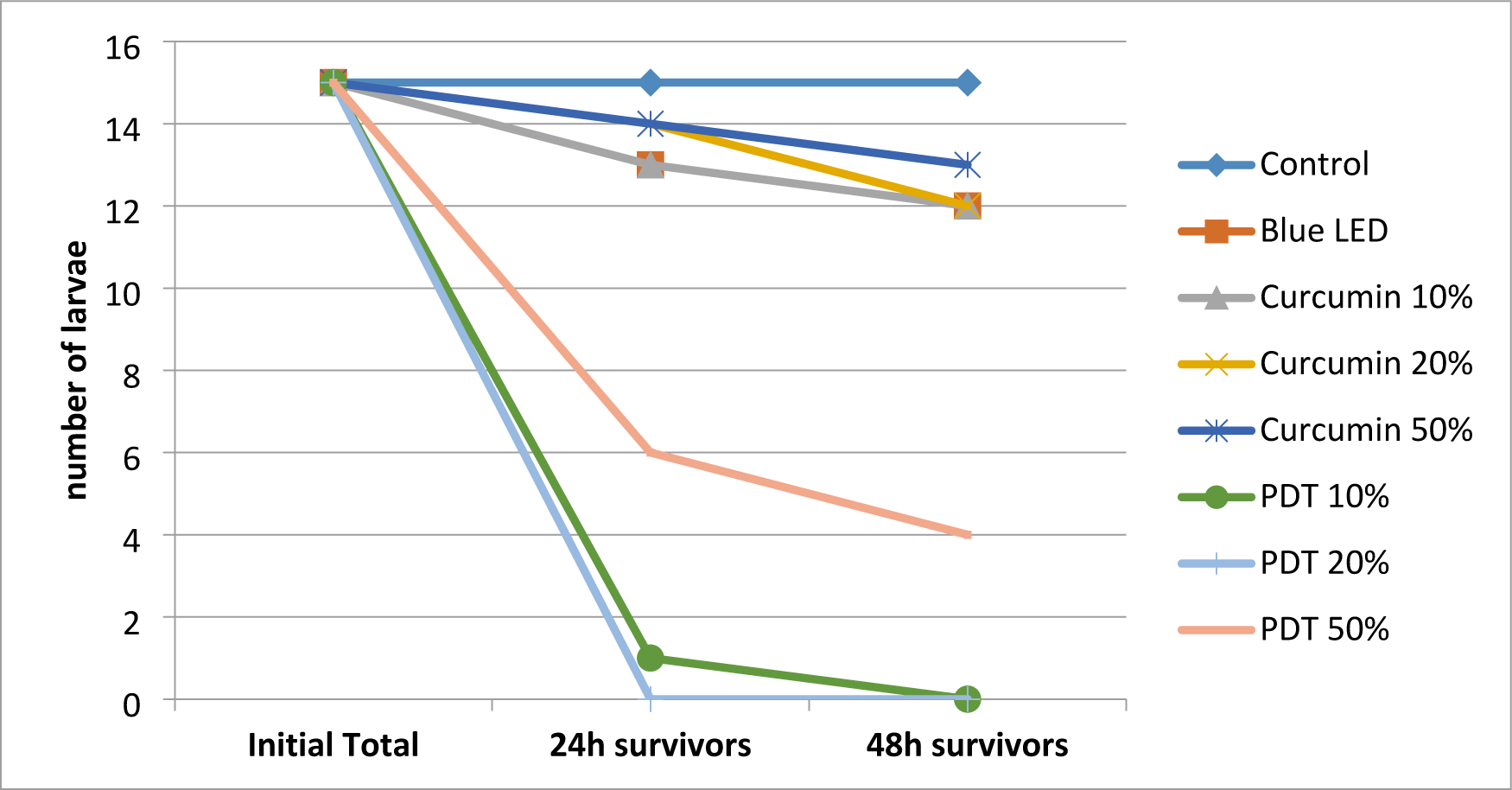
Survival of larvae after treatments in June / 2019 (test 1)

**Fig. 2.**
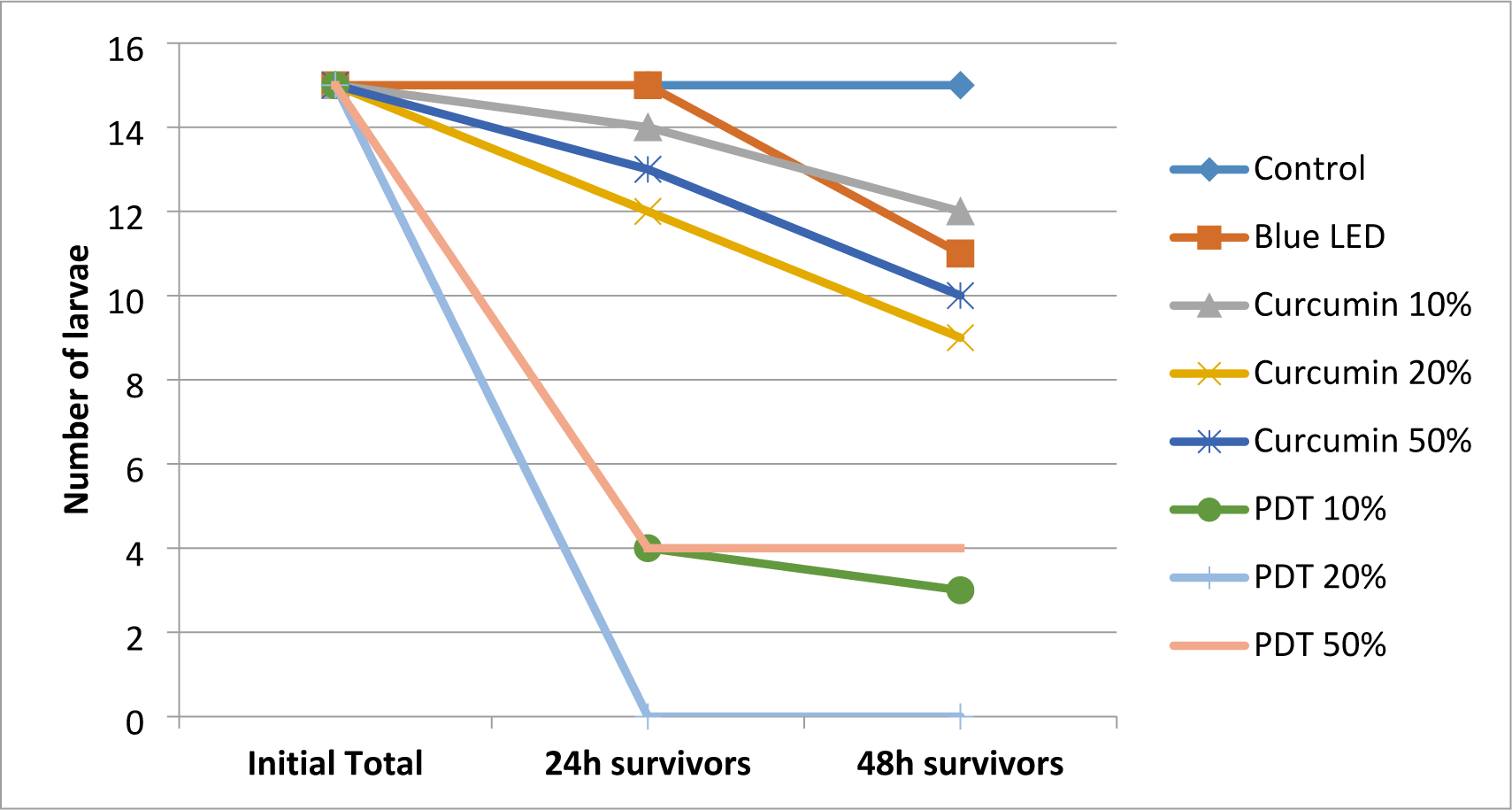
Survival of larvae after treatments in September / 2019 (test 2).

The effectiveness of technique given in percentage of larvae mortality in the studied groups was compared between the 24h and 48h times and it was observed in both test 1 and test 2, except for PDT 20% group (P + L + / 20%), that after 48h there is an increase in percentage of dead larvae in relation to the percentage of 24h. Only in test 2, PDT 50% group (P + L + / 50%) did not show a percentage increase in dead larvae within 48 hours. In PDT 20% group (P + L + / 20%), within 24 hours, 100% of tested larvae died, with no need to compare the percentage with 48 hours, since the total elimination occurred before that time, denoting the protocol’s efficiency, as shown in figures 3 and 4.

**Fig. 3.**
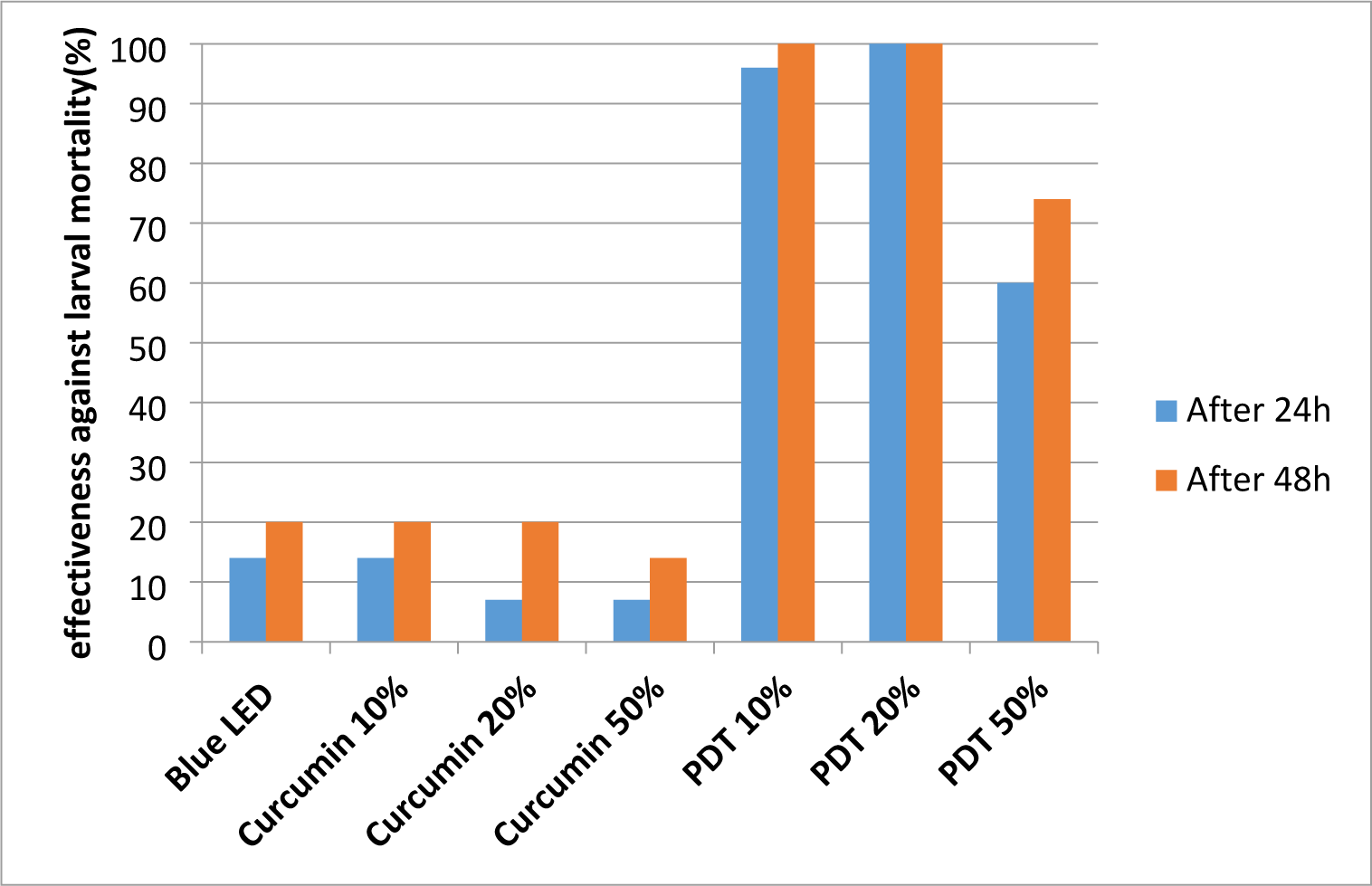
Effectiveness of techniques regarding larval mortality in June / 2019 (test 1)

**Fig. 4.**
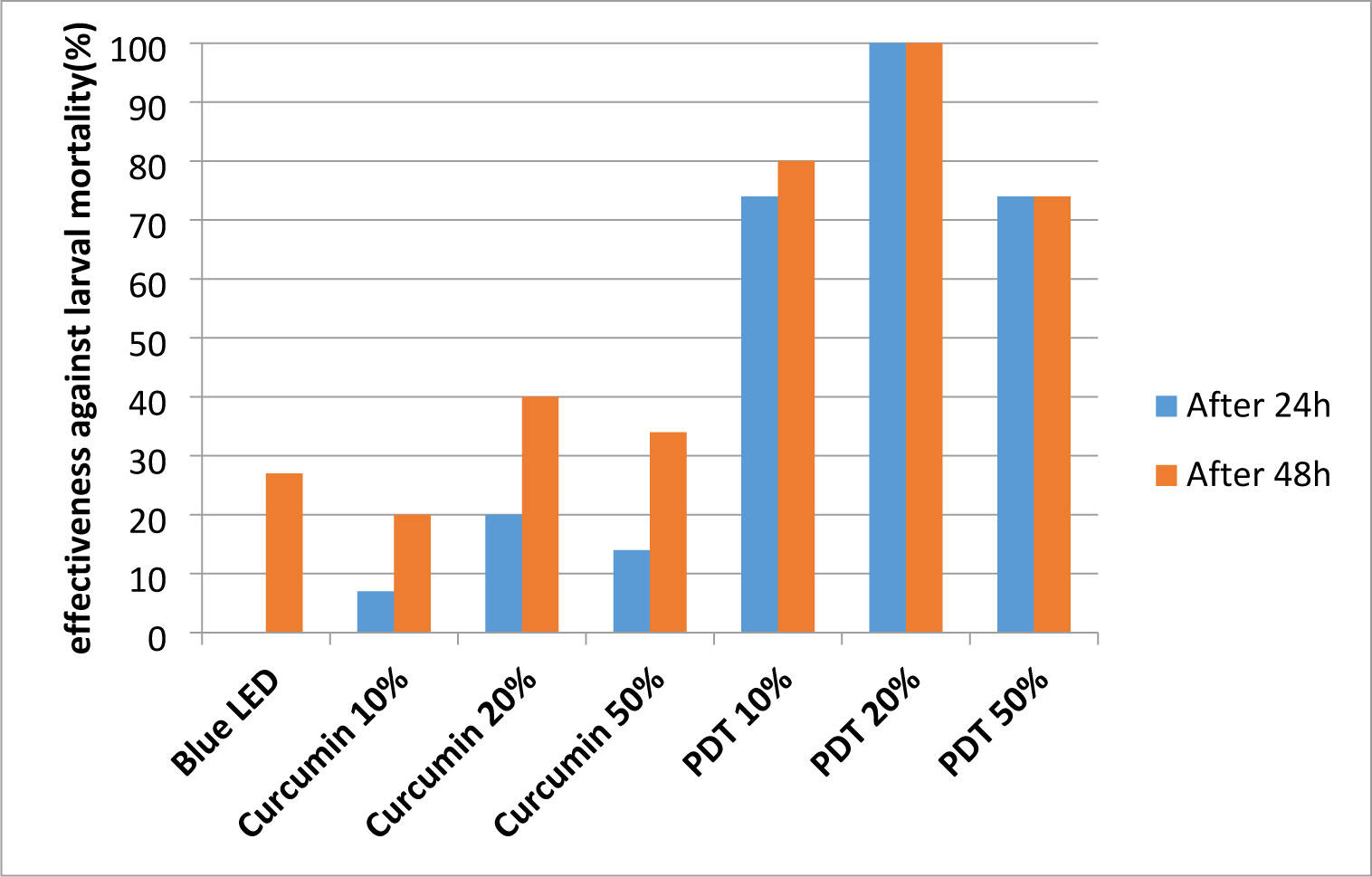
Effectiveness of techniques regarding larval mortality in September / 2019 (test 2).

### Histopathological Analysis

The larvae that were submitted to PDT according to parameters established for PDT 20% group (P + L + / 20%) presented a wide internal structural disorganization in relation to control larvae. In addition, they showed diffuse hypocellularity, a large number of cells with nuclear pycnosis and remaining cells with large vacuolization. In larvae that were subjected only to irradiation with blue LED and only fed with ration containing curcumin in the proportion of 20%, the presence of cells showing vacuolization and nuclear pycnosis is also observed, although in lesser number when compared to those which submitted to PDT, as shown in figure 5.

**Fig. 5.**
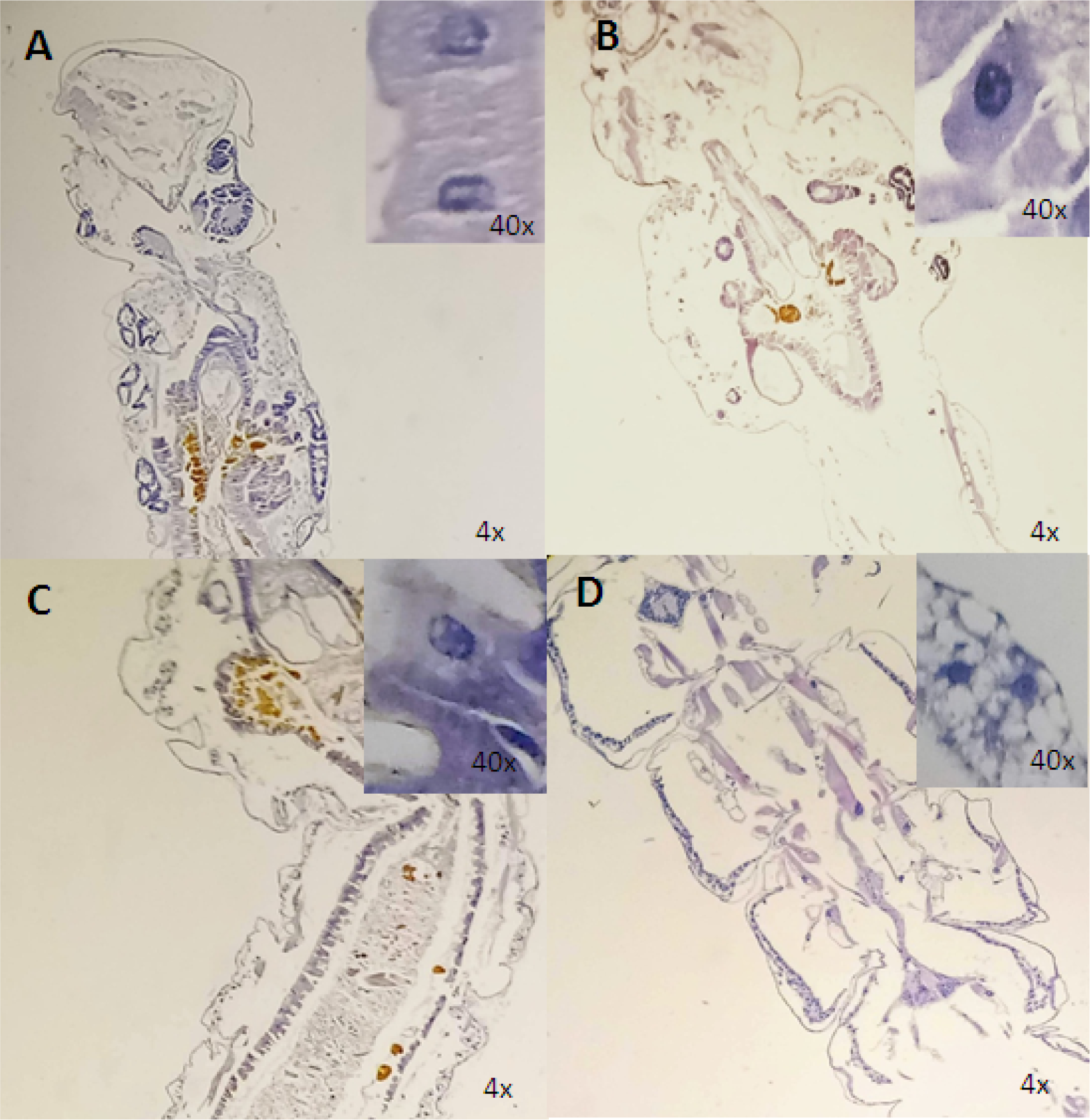
Histological sections of larvae belonging to control group (P-L-) (A); to blue LED group (P-L +) (B); to curcumin group 20% (P + L- / 20%) (C) and to PDT group 20% (P + L + / 20%) (D). There is a greater amount of vacuolated cells and a high degree of tissue destruction in larva treated with PDT compared to other treatments.

### Confocal Laser Scanning Microscopy

Confocal microscopy images showed larvae previously fed with ration mixed with photosensitizer curcumin in proportion of 20%, analyzed with excitation at 458nm and control larva, that is, that it wasn’t fed with the mixture containing curcumin. It was observed that the larvae had not presented 458 nm autofluorescence emission and in treatment images, apparently there was curcumin in both digestive and respiratory tracts, as shown in Figure 6.

**Fig. 6.**
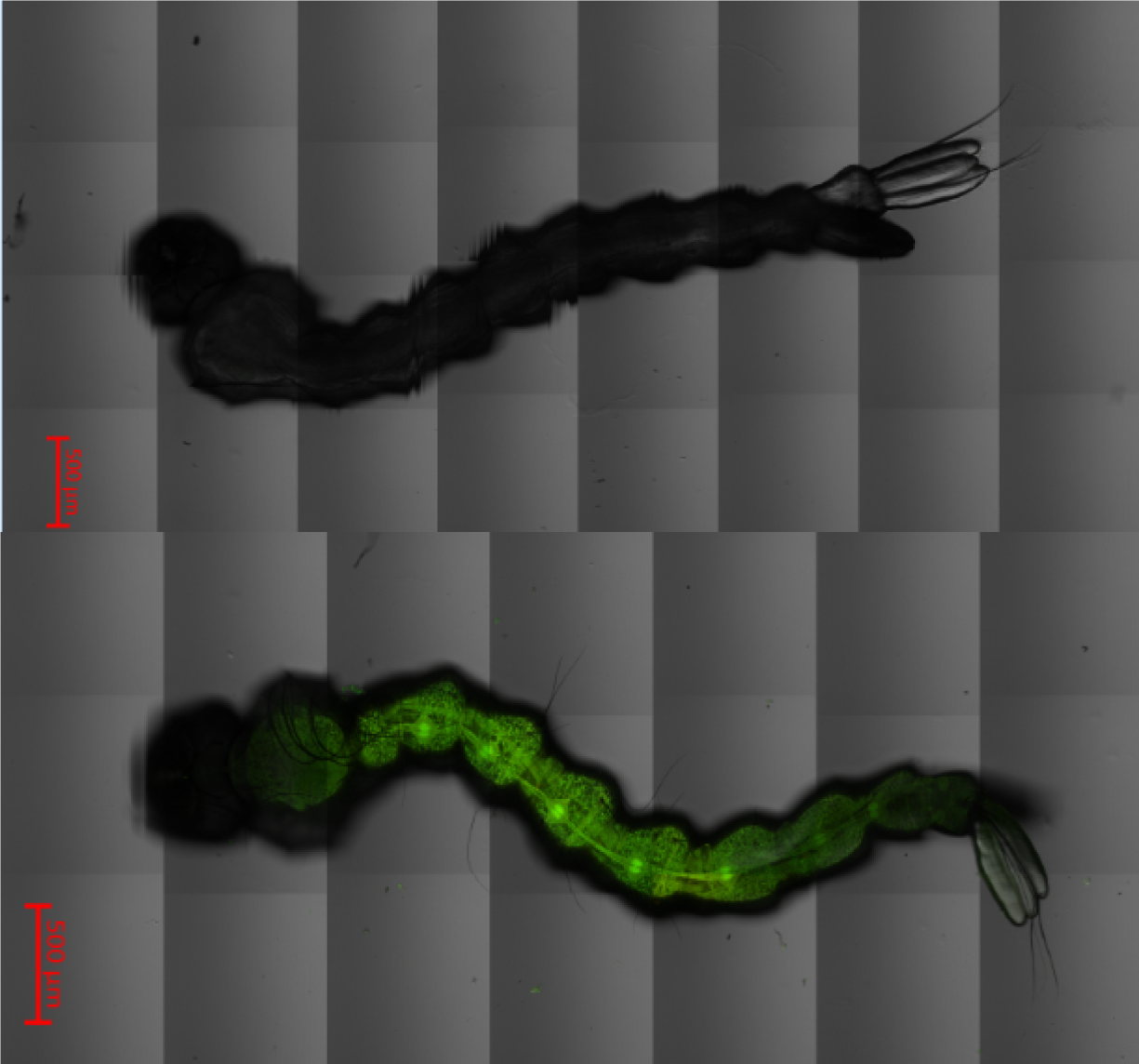
Images obtained by confocal microscopy of larva previously fed only with fish food and of larva fed with diet containing photosensitizer curcumin in proportion of 20%, where the presence of that compound is observed throughout the digestive tract.

### Raman microspectroscopy

Raman microspectroscopy of larval digestive tract was performed to analyze the molecular mechanisms of curcumin’s action as a photosensitizer. Figure 7 shows Raman spectra of the larva’s intestinal tissues, which did not feed of the diet containing photosensitizer curcumin at 20%, after the larva ingested the diet containing curcumin at a proportion of 20% and the same after exposure to LED light after feeding the mixture. The arrows indicate all Raman peaks that appeared or disappeared after the larva ingested the mixture as food or after exposure to LED. The 980 cm^-1^ peak, which appears in control spectrum and may belong to amino acid tryptophan, disappears or changes to 968 cm^-1^ after exposure to LED. Peaks of 1186 and 1244 cm^-1^ which may belong to nucleic acid guanine and Amide III protein Raman band, respectively, appear after the larvae are exposed to LED. The fluorescence profile (background signal in spectra) also changed and increased significantly after larvae ingested the mixture with 20% curcumin and especially after exposure to LED. All of those changes in Raman spectra and fluorescence mean changes in cells of larvae intestines, which can include denaturation of cellular proteins and cell death.

**Fig. 7.**
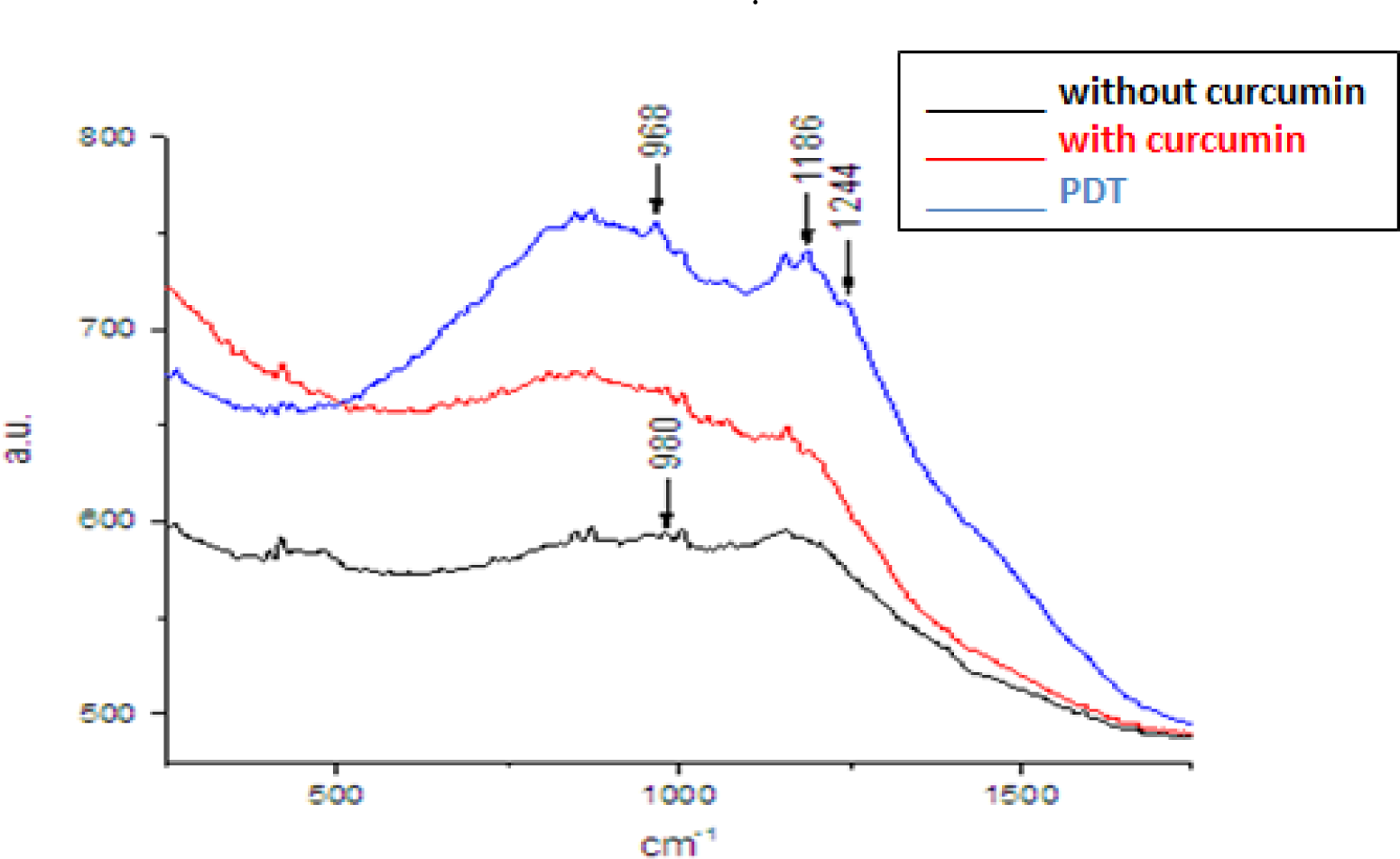
Raman spectra of larva intestine that did not ingest curcumin, after ingesting the fish ration and curcumin mixture 20% and exposure to LED after ingestion of the mixture. The arrows indicate peaks in spectra that appeared or disappeared after exposure to LED.

## DISCUSSION

The present study showed that photodynamic therapy is a technique with potential to be used effectively as part of strategies for vector control. Mortality of a small number of larvae was observed after treatments with only blue LED and only with food mixed with the photosensitizer in different concentrations, whereas the interaction of photosensitizer with light provided mortality of a greater amount of larvae. The photosensitizer curcumin, derived from saffron, does not cause any environmental or human health damage and, therefore, has a great advantage over substances widely used as larvicides [10]. The results showed that, when using the photosensitizer in proportion of 20% as a source of food for the larvae for the period of 24 hours and, later, irradiation with blue LED for at least 2 hours, the PDT proved to be a technique 100 % effective in eliminating the larvae within 24 hours after treatment.

Dondji et al. (2005) [11] evaluated the mortality of fourth stage larvae of *Aedes aegypti, Anopheles stephensi* and *Culex quinquefasciatus* induced by PDT mediated by photosensitizers bengal rose, chlorine e6, chloroquine, tetrafenil prophylamine porphyrin, tetrafenyl porphyrin and tetrahydrate, activated by 500 W tungsten lamps. In 72 hours of exposure to light, a mortality of 100%, 80% and 70% was observed, corresponding to use of bengal rose, hematoporphyrin and chlorine e6, respectively, which indicates that PDT is efficient also in the control of larvae of other species of mosquitoes that transmit diseases.

Lucantoni et al. (2011) [12] have studied the cationic porphyrin C14, incorporated into rodent feed, activated by fluorescent lamps from 400 to 800nm against *Aedes aegypti* larvae. Those conditions have showed high efficiency in photoinactivation of those organisms, leading to a mortality of 92%, after a period of 3 hours of illumination, corroborating with the results of current study.

The species *Aedes albopictus*, widely distributed in Brazilian territory and also responsible for transmission of arboviruses, also seems to be susceptible to alternative forms of vector control. Rodrigues et al. (2019) [13] have shown the larvicidal effect through enzymatic inhibition of extract of *Annona muricata* on larvae of species. The present study demonstrated the effectiveness of photodynamic therapy against mosquito larvae of the same genus.

Studies with *Curcuma longa* initially used only as a larvicide can be found. Sagnou et al. (2012) [14] have evaluated the larvicidal effectiveness of three curcuminoid pigments (curcumin, demetoxicurcumin and bisdemetoxicurcumin) against *Culex pippiens* larvae. The results have shown high larvicidal activity in larvae treated with curcumin with the mixture of three pigments. The larvae of mosquitoes of genus *Aedes*, unlike the species mentioned in this research, seem to be less sensitive to the isolated use of curcumin. However, the images obtained by laser scanning confocal microscopy in the current study showed that the photosensitizer was distributed throughout the digestive system of larva after being ingested, which certainly favored the large number of damages caused by PDT in the internal structure of larvae.

Essential oils and natural derivatives of *C. longa* were evaluated as possible larvicides by Anstrom and collaborators (2012) [15] and by Kalaivani and collaborators (2012) [9] in *Aedes aegypti* larvae, and the authors demonstrated that larvae on first and fourth stage were sensitive to the analyzed substances, showing their effectiveness as larvicides. However, the present study showed that the use of curcumin alone in three different proportions tested, in the same way as the isolated use of blue LED, did not show results similar to PDT in terms of larval mortality, inducing the death of a smaller number of individuals. Histological analyzes showed that isolated uses of blue LED and curcumin were related to cellular and structural maintenance of digestive system, however PDT induced structural changes, cellular stress and death troughout the all internal larvae systems. These findings suggests the existence of a synergistic effect in association of photosensitizer with light, reflected also in number of dead larvae in analyzed times after the application of tests.

The use of essential oils to control *Aedes aegypti* is well studied and the results demonstrate that there is possibility of a series of effective alternatives for this purpose. Barreto and collaborators (2006) [16] subjected *Aedes aegypti* larvae to the crude ethanolic extract of *Sapindus saponaria Lin* (Sapindaceae), whose effectiveness as a possible alternative larvicide was demonstrated by a morphological study that showed damage to cells of digestive system of larvae. In current study, such damage was also obtained with PDT, as demonstrated by Raman microspectroscopy analysis, also showed cell damage and death. Histological sections showed intense cytoplasmic vacuolization in cells remaining after PDT, with a high degree of tissue destruction and loss of larva’s internal structure.

Costa et al. (2012) [17] have tested the extract of *Annona Coriacea* (Magnoliales: Annonaceae) to eliminate *Aedes aegypti* larvae and its activity was demonstrated through histological analysis of their digestive tract. The tested extract has caused cytoplasmic vacuolization of regenerative and columnar cells, leading the cells to apoptosis. A similar result was obtained by the present study, whose histopathological analysis showed that PDT caused diffuse hypocellularity, a large number of cells with nuclear pycnosis and remaining cells with great vacuolization.

The use of other synthetic substances, different from traditional larvicides, was also studied. Farnesi et al. (2012) [18] tested the use of a synthetic inhibitor of chitin synthesis called Novaluron in *Aedes aegypti* larvae. Morphological analyzes have shown that the substance had induced the formation of a discontinuous and altered cuticle at several points with the epidermis showing itself as thinner or absent. The essential oil of Schinus terebinthifolia has also caused changes in the deposition of chitin in epidermis of *Aedes aegypti* larvae, in a study by Pratti et al. (2015) [19]. The analysis by laser scanning confocal microscopy, in the present work, showed that curcumin is easily distributed among various larval tissues and further studies can be performed in order to verify its possible epidermal action.

Substances isolated from essential oils are also evaluated as alternatives for vector control. Valotto et al. (2014) [20] have shown that 3-β-acetoxylabdan-8 (17) -13-dien-15-oic acid, isolated from the medicinal plant *Copaifera reticulata* (Leguminosae), had caused morphological changes in *Aedes aegypti* larvae, as well as PDT in the present study.

As a form of biological control, the use of bacteria of genus *Wolbachia*, which are responsible for preventing the transmission of viruses when present in wild populations of mosquitoes, is widely studied. Silva et al. (2008) [21] have evaluated the action of bacterium *Bacillus thuringiensis var. israelensis* in *Aedes albopictus* larvae. Histological analyzes have shown that intestinal cells had presented disorganization of basal processes, dilation and fragmentation of rough endoplasmic reticulum, with intense cytoplasmic vacuolization after infection with the bacteria, demonstrating important changes such as those obtained by PDT in the present study. Thus, vector control alternatives that are not aggressive to environment and human health must be adapted for large-scale use to reduce the damage caused by indiscriminate use of insecticides over the past decades. PDT, in addition to proving to be effective, does not induce the appearance of any resistance mechanism, which represents a great advantage for its use.

## CONCLUSION

Photodynamic therapy was effective in controlling larvae of mosquitoes of genus *Aedes* in all tested parameters, with emphasis on group PDT 20% (P + L + / 20%), which has eliminated 100% of the larvae after 24h. Analyzes by confocal laser scanning microscopy showed that photosensitizer curcumin was distributed throughout the digestive system of larvae, facilitating the action of PDT, which caused cell damage and tissue destruction, which possibly culminated in the death of larvae.

## ACKNOWLEDGMENTS

We thank to PPSUS SUS0037/2018 and CAPES for supporting our research.

